# Deep learning from videography as a tool for measuring *E. coli* infection in poultry

**DOI:** 10.1101/2024.11.20.624075

**Authors:** Neil Scheidwasser, Louise Ladefoged Poulsen, Prince Ravi Leow, Mark Poulsen Khurana, Maider Iglesias-Carrasco, Daniel Joseph Laydon, Christl Ann Donnelly, Anders Miki Bojesen, Samir Bhatt, David Alejandro Duchêne

## Abstract

Poultry farming is threatened by regular outbreaks of *Escherichia coli* (*E. coli*) that lead to significant economic losses and public health risks. However, traditional surveillance methods often lack sensitivity and scalability. Early detection of infected poultry using minimally invasive procedures is thus essential for preventing epidemics. To that end, we leverage recent advancements in computer vision, employing deep learning-based tracking to detect behavioural changes associated with *E. coli* infection in a case-control trial comprising two groups of 20 broiler chickens: (1) a healthy control group and (2) a group infected with a pathogenic *E. coli* field strain from the poultry industry. More specifically, kinematic features derived from deep learning-based tracking data revealed markedly reduced activity in the challenged group compared to the negative control. These findings were validated by lower mean optical flow in the infected flock, suggesting reduced movement and activity, and post-mortem physiological markers of inflammation which confirmed the severity of infection in the challenged group. Overall, this study demonstrates that deep learning-based tracking offers a promising solution for real-time monitoring and early infection detection in poultry farming, with the potential to help reduce economic losses and mitigate public health risks associated with infectious disease outbreaks in poultry.

## 1. Introduction

Poultry farming is a cornerstone of the agriculture industry and is essential to global nutritional security, accounting for nearly 35% of the world’s meat production [1]. In many poultry-producing regions, *Escherichia coli* (*E. coli*) infection is a leading cause of morbidity and mortality, including China [2, 3], the United States [4], Brazil [5], and several European countries such as the United Kingdom [6] and Denmark [7]. In addition, *E. coli* outbreaks are linked with intensive use of antibiotics, promoting the selection of resistant strains that have the potential to spill over into human populations [8]. Consequently, detecting early signs of *E. coli* infection in domesticated birds, such as broiler chickens, is crucial for safeguarding animal welfare, minimizing economic losses, and protecting public health.

Over recent decades, a plethora of techniques have been proposed to leverage data from physiological markers (e.g., body temperature [9, 10]) or behavioural outputs (e.g., locomotion, sound [11]) to assess welfare impairments and detect pathogen outbreaks. Measuring temperature requires precise thermal imaging cameras, which are expensive and sensitive to environmental disturbances. In contrast, acquiring sound and video recordings is logistically simpler and less expensive, but usually requires computer-intensive postprocessing to extract meaningful behavioural features. In addition, obtaining precise estimates of individual patterns remains challenging. For instance, commonly used video processing methods such as optical flow have been leveraged to provide valuable insights on behaviour [12, 13], bacterial infection [14, 15], footpad dermatitis and hockburn [16], and general animal welfare [17–19]. However, optical flow only allows for analysis at the level of a flock and not of the individual. Additionally, as it relies on the average displacement of pixels across consecutive frames, it is therefore prone to capturing irrelevant movements from the environment. To obtain individual-level measurements, gait-scoring systems have been developed for broiler chickens [20, 21], but are impractical for real-time applications. Other frameworks based on markers or wearable sensors (e.g., accelerometers or radio-frequency identification (RFID) chips [22]) are promising for their high resolution, but are intrusive and not easily scalable to industrial settings.

Alternatively, recent progress in deep learning [23] for computer vision has catalysed the development of versatile frameworks for tracking and markerless pose estimation (i.e., without pre-placed physical markers) of individual [24–26] or multiple animals simultaneously [27–29]. These methods have garnered significant interest in behavioural neuroscience [30] and ecology [31]. As to animal behaviour and welfare monitoring, the potential advantages of deep learning-based videography analysis are multi-factorial. Indeed, state-of-the-art solutions are non-invasive, adaptable to various environments, user-friendly, and may allow for individual-level analysis with near-human-level precision. Although these systems initially require data-intensive training of deep neural networks on videos of interest, transfer learning allows for fast deployment of pre-trained models on new video data.

To this end, we develop a feasibility study that demonstrates the capacity of deep learning-based, markerless tracking from video recordings to detect phenotypical and physiological differences in broiler chickens following *E. coli* infection. We designed a trial in which 40 chickens were randomly assigned to two, equal-sized, separately-housed groups: (1) a healthy control group (control), and (2) a group infected with a pathogenic *E. coli* strain (Fig. 1). We used the multi-animal version of DeepLabCut [28] to identify behavioural changes associated with infection and found that *E. coli*-infected chickens moved and travelled less in their pens and spent less time near the food source (used as a proxy to measure feeding time). This finding was first validated with a more traditional optical flow-based analysis, which corroborated the lower activity in the infected group, and second with necropsy data which revealed distinct patterns of *E. coli*-caused internal lesions and inflammation in the infected group. Thus, our data provide evidence that deep learning on video is a useful and objective tool for detecting the behavioural and physiological changes associated with infection. The simplicity and objectivity of the statistics gleaned from videos, including distance travelled and changes in body area, indicate that scalability to industrial settings is feasible. Addressing sensitivity to other pathogens, high animal densities, and trade-offs between labelling individual animals and image resolution will be crucial for implementing reliable deep learning-based outbreak detection methods in broiler farms.

**Figure 1.**
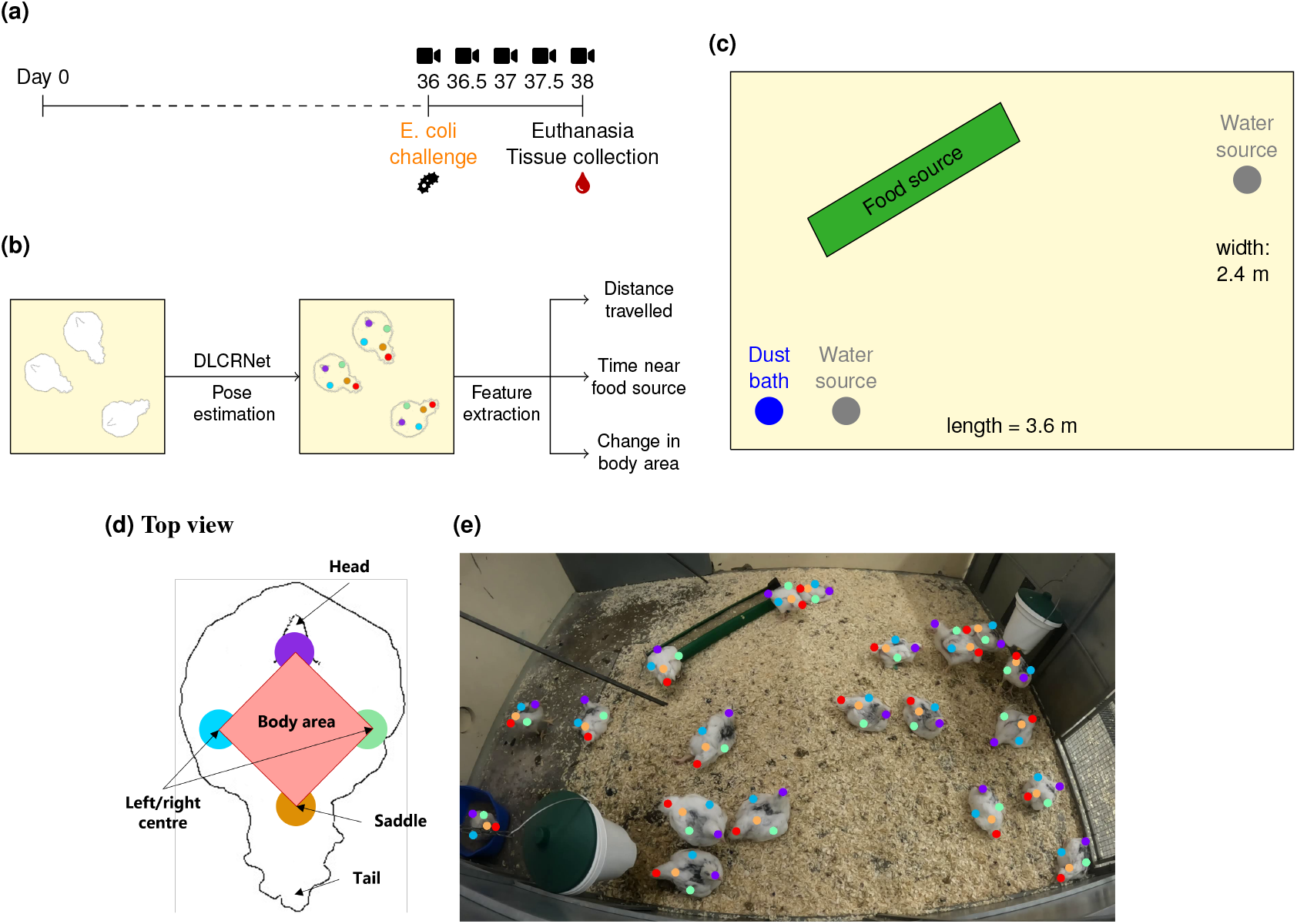
Experimental setup. **(a)** Timeline of the experiment. **(b)** Pipeline for behavioural feature extraction using DLCRNet, the standard model of the multi-animal version of DeepLabCut (maDLC) [28]. **(c)** Diagram of the chicken coops. **(d)** Standardised body parts used for tracking and polygon used to approximate body area. **(e)** Example predictions from maDLC, based on one frame.

## 2. Methods

### 2.1. Animals and housing conditions

40 day-old, *E. coli*-unvaccinated, Ross 308 broiler chickens (mixed gender) were obtained from Blenta AB (Blentarp, Sweden) and followed for 38 days (Fig. 1a). The animals were accommodated in two groups of 20 (see Experimental design) in an 8.64 m^2^ coop (3.6 m x 2.4 m; Fig. 1c) with a dust bath to a 16h light (06h-22h):1h dim (22.00-23.00):7h dark (23.00-06.00) schedule. Feed was provided *ad libitum* twice daily to accommodate the increased growth (from 15 to 200 grams/day) according to the feeding schedule of Aviagen [32]. The feed consisted of commercial whole-grain food for broiler chickens aged 0-8 weeks (Brogaarden, Lynge, Denmark). Water was available *ad libitum* at two different sources. Standard procedures were conducted to prevent cross-contamination from other pathogens.

### 2.2. Experimental design

On the day of arrival, each chicken was randomly assigned to one of two groups: uninfected control group, or infected with *E. coli*. For the infected group, a pathogenic strain of *E. coli* known for causing colibacillosis in broilers [33–35] (strain ST117 E44; accession number LXWV00000000.1) was administered intratracheally (9.9 × 10^5^ colony-forming units (CFUs)/chicken in 0.5 ml) (week 5 of the experiment; Fig. 1a), while chickens from the control group were sham-challenged intratracheally (phosphate-buffered saline; (PBS)) following the same timeline. The strain used for inoculation was provided by the Section of Veterinary Clinical Microbiology at the University of Copenhagen and prepared as described in [35] (storage at -80°C, streaking on blood agar base supplemented with 5% bovine blood (BA) and incubation at 37°C overnight). A single colony was incubated in brain heart infusion (BHI) broth (Oxoid BO1230B (Thermo Fisher Scientific)) overnight at 37°C. Subsequently, 1 ml of the culture was transferred to 100 ml BHI and incubated until the optical density at 600 nm (*OD*_600_) reached 1.56 (approximately two hours). At *OD*_600_ = 1.56 (exponential phase), 308 µL of culture was transferred to 199.69 ml PBS (Merck, 524650) and a volume of 0.5 ml per animal was used as inoculum and kept on ice until infection. The inoculum’s CFU count was confirmed by 10-fold dilutions prepared in triplicates plated (100 µL) on BA, and incubated overnight before counting.

### 2.3. Biological data sampling

Blood samples were collected twice from the brachial vein: firstly a day before the challenge and secondly before euthanisation for analysis of serum amyloid A using an LZ-SAA assay (Eiken Chemical). Blood samples were centrifuged at 1,500 RPM for 10 min to isolate the serum, which was thereafter stored in a freezer (−20°C) until analysis. Euthanisation was performed by cervical dislocation directly following the induction of unconsciousness by blunt head trauma. Gross pathology was performed two days post-infection during the necropsy in a randomised and blinded manner to mitigate experimenter biases. An overall lesion score (see Table S1) derived from previous *E. coli studies* [35–37] was calculated by visually assessing and grading pathological lesions in the peritoneum, air sacs, lungs and spleen. To measure pulmonary *E. coli* CFU counts, each lung was aseptically removed and placed in a sterile stomacher bag, weighed, and then blended with a 1/1 (weight/vol) solution of 0.9% PBS. The resulting blend was serially diluted tenfold (10^*−*1^-10^*−*7^) and 3 × 30 µL per dilution was spotted separately for overnight incubation. CFU counts were normalised by lung weight.

### 2.4. Behavioural data sampling

For each group, five videos were recorded over three days: two in the first two days (morning and afternoon, after feeding) and one in the third morning (Fig. 1a). Each recording lasted between 20 and 30 minutes and was acquired at 30 frames per second (FPS) at a resolution of 1080 × 1920 pixels using GoPro HERO10 Black cameras (https://gopro.com). The cameras were placed 130 cm away from the floor. For each video, the first two minutes were removed to minimise any potential experimenter intervention.

### 2.5. DeepLabCut training and inference

Deep learning for animal tracking was implemented in markerless fashion using the multi-animal version of DeepLabCut [28], which allows tracking and pose estimation of multiple objects simultaneously. For each video, each frame from a single video is first clustered using *k*-means clustering, and twenty frames were sampled from the formed clusters (100 frames is considered sufficient to achieve reasonable tracking performance according to [24]). For each frame, annotations for five body parts (head, centre left, centre right, saddle, and tail) were added for each animal. We used a pre-trained *DLCRNet_ms5* [28], which consists of a multi-scale convolutional neural network (CNN) stacked on top of another CNN (ResNet-50 [38]) for feature extraction and stacked with deconvolutional layers to predict where the body parts of interest were and which animals they belonged to. The multi-scale architecture fuses high and low-resolution maps, thus allowing for robust and accurate detection of body parts by leveraging fine-grained detail and broader contextual information. The model was fine-tuned in end-to-end fashion (i.e., all model weights were tunable) with recommended hyperparameters from [28] (Adam [39] optimisation with multi-stage learning rate (*η*) scheduling (stage 1: *η* = 1 × 10^−4^ for 7,500 iterations; stage 2: *η* = 5 × 10^−5^ for 12,000 iterations; stage 3: *η* = 1 × 10^−5^ until the end of the experiment), batch size of 8) using 95% of the labelled frames (split randomly) for 200,000 training iterations. The root mean squared errors (RMSEs) were: 3.03 pixels (train set) and 8.68 pixels (test set) for an image size of 1080 × 1920 pixels. The trained network was then used to extract the (x, y) positions of each body part for each animal in all videos. An example frame is shown in Figure 1e.

### 2.6. Post-processing

For each video, DeepLabCut outputs an array of (x-coordinate, y-coordinate, and likelihood) triplets of size *F* × (*N × B* × 3), where *F* is the number of frames, *N* the number of animals, and *B* the number of body parts. Predictions with a likelihood below 0.9 were dropped following standard DeepLabCut practice [24]. To recover some of the dropped values, forward linear interpolation was applied at a body part level, where at most 15 consecutive missing values could be filled (0.5 s). (x, y) positions were finally smoothed using Savitzky-Golay filtering with a polynomial order of 7 and a window length of 15 frames to strike a trade-off between preserving the sharpness of the original signal and reducing noise. Subsequently, two kinematic features could be extracted (Fig. 1b) for each chicken *c* in video *v*: the total distance travelled *L*_.,*v,c*_ to quantify locomotor activity and the total rate of change in body area 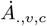, for non-locomotor activity (e.g., stretching for improved ventilation).

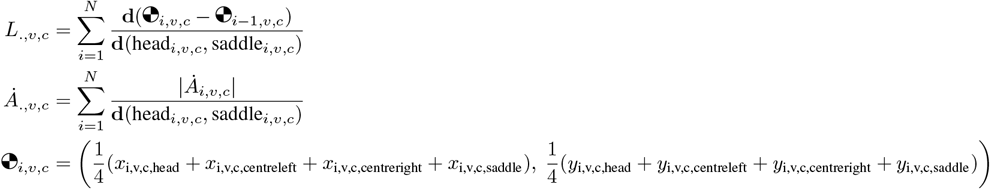

The distance travelled in each frame *i* is calculated as the Euclidean distance **d** between the chicken’s centre of mass 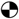 in the current frame and the previous frame *i* − 1. The centre of mass is defined as the average position of four key points: the head, centre-left, centre-right, and saddle. For each frame *i*, both *L*_*i,v,c*_ and 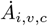 are normalised by the distance between the head and saddle in each respective frame to account for potential radial and tangential distortions due to the camera setup. The frame-wise rate of change in body area was measured in a similar fashion to [40]. The shoelace formula was employed to calculate the area of a polygon defined by the four above-mentioned key points (Fig. 1d). The rate of change of that area 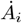 was measured as a discrete temporal derivative of the body area, obtained numerically using a Savitzky-Golay filter. Subsequently, the time spent near the food source (assumed as a proxy for time spent eating) was estimated for each animal by computing the number of frames in which the animal’s head was in the vicinity of the food source (Fig. 1c). The vicinity was defined as a rectangle whose scale was 1.05 times that of the food source in the video. Future work should investigate more direct measures of feeding behaviour, e.g., by detecting pecking or other related actions, to increase the accuracy of this measurement.

### 2.7. Statistical analysis

For each physiological covariate, individual differences were tested using non-parametric Mann-Whitney U tests. The *t*-test assumption of normality was assessed using quantile-quantile plots (Q-Q plots), while Levene’s test was used to test for homoscedasticity. For all markers, both normality and homoscedasticity could not be reasonably assumed for lesion scores and SAA levels, leading us to opt for Mann-Whitney U tests to assess between-group disparities. Cohen’s *d* was reported to reflect the effect size for all pairwise comparisons.

For each behavioural feature, a Bayesian mixed-effects model was employed to investigate between-group variations (with the control group as a reference) while accounting for random variations across the different recordings. Our mixed effects model had the following linear predictor

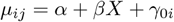

where *µ*_*ij*_ is a behaviour feature of interest (e.g., distance travelled) for a day *i* and individual *j, α* is an intercept that corresponds to the control group, *β* is a scalar for the group *X* (infected or not), and *γ* is a random effect for the *i*^*th*^ day. As the distance travelled and body area change were normalised by the body length (thus constituting rates distributed in (0, 1)), we assumed their values were generated from a Beta distribution. Whilst the time spent near the food source is also a rate, a zero-inflated version of the Beta distribution was used to account for the frequent zero-valued observations. All models were fitted with the default prior distributions suggested by `brms` [41]. To facilitate efficient sampling and minimise divergent transitions, a target acceptance rate of 0.99 was chosen and a maximum tree depth of 15 was selected for building during the trajectory phase of the Hamiltonian Monte Carlo sampler. For all variables, we compared the beta family models (with a logit link function) against models with the default family parameter (Gaussian with an identity link function) in `brms`. We selected the best family based on a Bayesian estimate of the expected log pointwise predictive density (ELPD) from leave-one-out cross-validation (LOOCV) (Table S4). Finally, convergence plots and posterior predictive checks were plotted to check model fit (see Fig. S1 to S4). For all variables, the models fitted using a beta distribution were preferable.

### 2.8. Optical flow

To complement deep learning-based animal tracking with more traditional computer vision methods, we measured optical flow, a commonly used technique to determine the average motion of chickens in a video as a means to assess animal welfare [12, 14, 42]. Optical flow works by estimating the motion of pixels in a video by comparing consecutive image frames. Here, the optical flow was computed using a state-of-the-art adaptation of the Recurrent All Pairs Field Transforms (RAFT) model [43] called SEA-RAFT [44]. This model consists of a deep neural network (DNN) trained to predict the optical flow between consecutive frames. It features a residual neural network (ResNet-34 [38]) to extract relevant features from pairs of images, a correlation block to compute similarities between pixels, and a recurrent neural network (RNN) to iteratively refine the flow field predictions from the correlation block outputs. Compared to traditional, handcrafted methods such as the Horn-Schunck [45] or Farnebäck methods [46], which are based on energy minimisation, deep learning-based approaches have been posited to be more robust to occlusions and rapid movement [47].

Here, we used a version of SEA-RAFT that was pre-trained on a collection of datasets including simulated moving objects, photorealistic scenes, and recorded scenes from autonomous driving vehicles [44]. To reduce inference time, we downsampled the images by a factor of 2 (*h* = 960 × *w* = 540 pixels) before feeding them into the model. For each pair of frames, the output consists of an array of size 2 × *h* × *w*, which denotes the estimated (*x, y*) flow for each pixel. From this output, following previous optical flow-based animal welfare studies [12, 14, 42], the first four moments (mean, variance, skewness, and kurtosis) of the optical flow norms were measured for each frame. The statistics were finally averaged across frames for between-group comparisons, and the average mean optical flow was regressed using ordinary least squares (OLS) against the distance travelled measured from DeepLabCut-based tracking.

### 2.9. Implementation

Video pre-processing was performed using `ffmpeg` (https://ffmpeg.org) and tracking was implemented using the DLCRNet model [28] implemented in `DeepLabCut` [24] (version 2.3.5). Model training on our video dataset was performed on a single NVIDIA RTX A6000 graphical processing unit (GPU). Optical flow was computed frame by frame using an implementation of a pre-trained SEA-RAFT model [44] in `PyTorch` [48] (version 2.2.0). Feature extraction and statistical tests were performed using custom scripts in Python 3.10.10 using `NumPy` [49] (version 1.25.0), `SciPy` [50] (version 1.11.1), and `Pingouin` [51] (version 0.5.4). Bayesian mixed-effects modelling was implemented in R 4.4.0 using the `brms` package [41] (version 2.21.0). The code for reproducing feature extraction and statistical analyses is available at https://github.com/Neclow/dlc4ecoli.

### 2.10. Ethics

The study was approved by the Danish Animal Experiments Inspectorate under the Danish Ministry of Environment and Food, and all animal procedures were performed by this approval (license no. 2019-15-0201-01611) and with ARRIVE guidelines. The licence granted, and guidelines hereof are in agreement with the Danish law on animal experiments and the EU directive 2010/63. Predefined humane endpoints were determined and animals were observed every 30 min for the initial six hours after inoculation, subsequently at maximally eight-hour intervals for three days after inoculation with increased frequency in the event of clinical signs in any of the groups. If clinical signs were present, e.g., ruffled feathers, depression, anorexia, lethargy or dyspnoea, the bird was either treated with 0.1 mg/kg buprenorphine and observed with increased frequency or euthanised.

## 3. Results

The goal of this study was to determine whether deep learning-based behavioural analysis from video can be used as a proxy for monitoring and detecting animal welfare impairments (here, caused by *Escherichia coli* (*E. coli*) infection). To this end, we recorded two groups of 20 animals: a negative control group and a group infected with *E. coli*. In particular, we compared standard physiological measurements obtained post-mortem (see Methods) against behavioural features extracted from multi-animal DeepLabCut [28].

Post-mortem examination of the chickens by necropsy revealed significant disparities between the groups in terms of lesion score, SAA levels, and number of colony-forming units (CFUs) (Fig. 2; Mann-Whitney U: *U*_*lesion*_ = 39.5, *U*_*SAA*_ = 45.0, *U*_*cfu*_ = 31.0; *p*-values *<* 0.001) with medium to large effect sizes (Cohen’s *d*: *d*_*lesion*_ = -2.121, *d*_*SAA*_ = -1.153, *d*_*cfu*_ = -0.607). Chickens from the control group had near-zero presence of SAA and CFUs as well as very few lesions, while *E. coli*-infected groups showed significantly higher values for all markers (Table S2).

**Figure 2.**
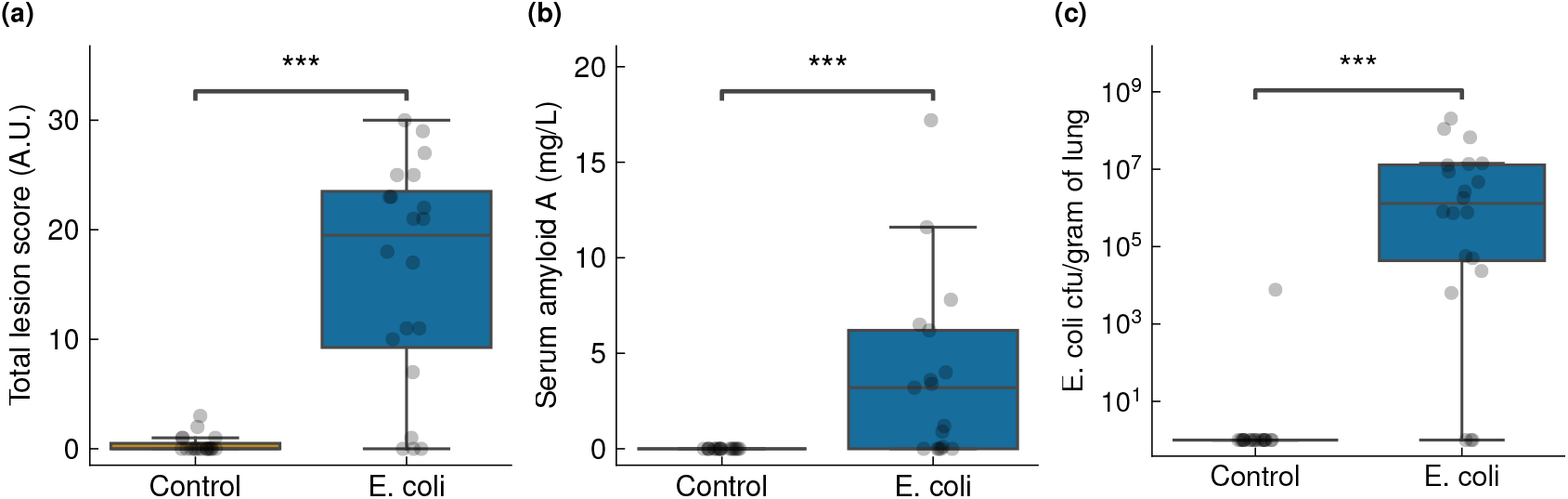
Comparison of physiological biomarkers of inflammation in response to *E. coli* challenge. Significance levels: \**P <* 0.05, \*\**P <* 0.01, \*\*\**P <* 0.001.

Behavioural differences between groups in the distance travelled, body area, and time near the food source (extracted from DeepLabCut-based animal tracking) were evaluated using Bayesian mixed-effects modelling, with random effects applied to the recording time (Fig. S1-S4). The infected group was found to be significantly less active than the uninfected control group as measured by the distance travelled (average treatment effect (ATE) ratio: 24% reduction (95% credible interval (CrI): 6-39%)) and the rate of change in body area (ATE ratio: 23% reduction (95% CrI: 4-39%)) (Fig. 3; Table S3). Additionally, we posit that the infected group fed less as its individuals spent less time near the food source on average (ATE ratio: 41% reduction (95% CrI: 24-56%); Fig. 3). Individual chicken identity was not recovered from one recording to another, such that a chicken labelled “chicken X” in a given recording could not be re-identified in subsequent recordings, preventing within-subject analyses.

**Figure 3.**
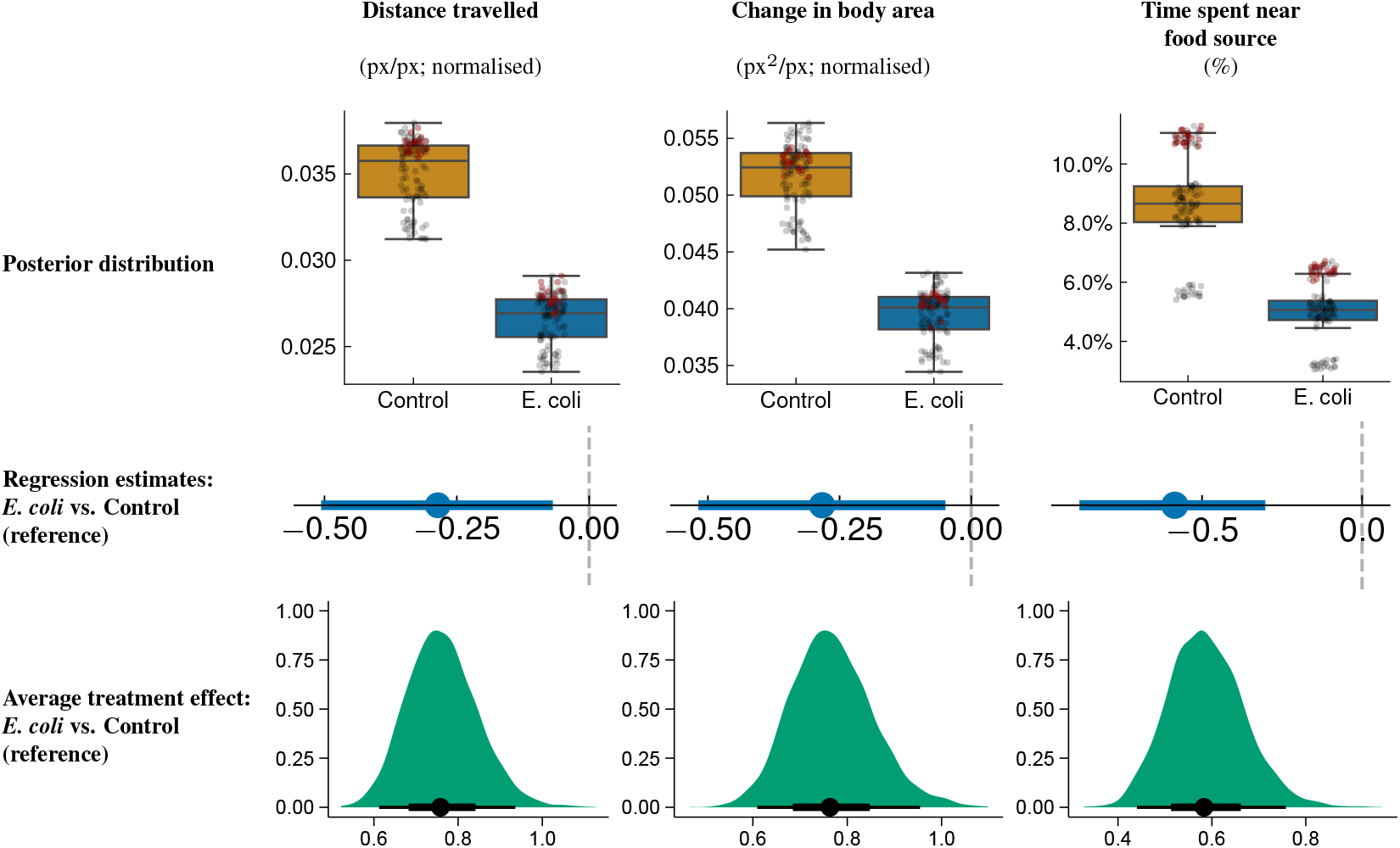
Comparison of behavioural features in response to *E. coli* challenge. Top: posterior distributions from Bayesian mixed-effects modelling to investigate between-group variations while accounting for random variations across the different recordings, predicting the group (infected or not). One point refers to one individual in one video. Points from the first recording (performed on challenge day) recording are marked in dark red. Middle: mean estimates of the same behaviour features for the *E. coli* group, with the negative control group serving as the reference. Bottom: average treatment effect for each behaviour feature, computed as a ratio between the *E. coli* and control groups. Error bars indicate 95% confidence intervals.

The activity measured by deep learning-based individual tracking was validated against optical flow, a traditional method in computer vision used to measure object motion based on pixel intensity changes between consecutive frames (Fig. 4). Optical flow data from videos (Control: *n* = 5, Infected: *n* = 5) of the control group exhibited higher mean optical flow (Fig. 4a), indicating higher average activity [14]. Whilst not exhibiting higher optical flow kurtosis as in [14], control individuals also displayed higher optical flow variance, indicating that infected individuals had more uniform behaviour. The optical flow results were generally consistent with DeepLabCut tracking-based results, seen in the positive correlation between the mean optical flow and the mean distance travelled (Fig. 4b to 4d; group- and time-independent Pearson correlation: *r* = 0.636, *p* = 0.048; *BF*_10_ = 2.17). Overall, despite the limited validation provided evidence by optical flow due to sample size limitations, deep learning-based video analysis could be deemed effective at discriminating behavioural patterns between the control and infected groups. Further, the impact of infection was confirmed by traditional post-mortem analysis of infection biomarkers.

**Figure 4.**
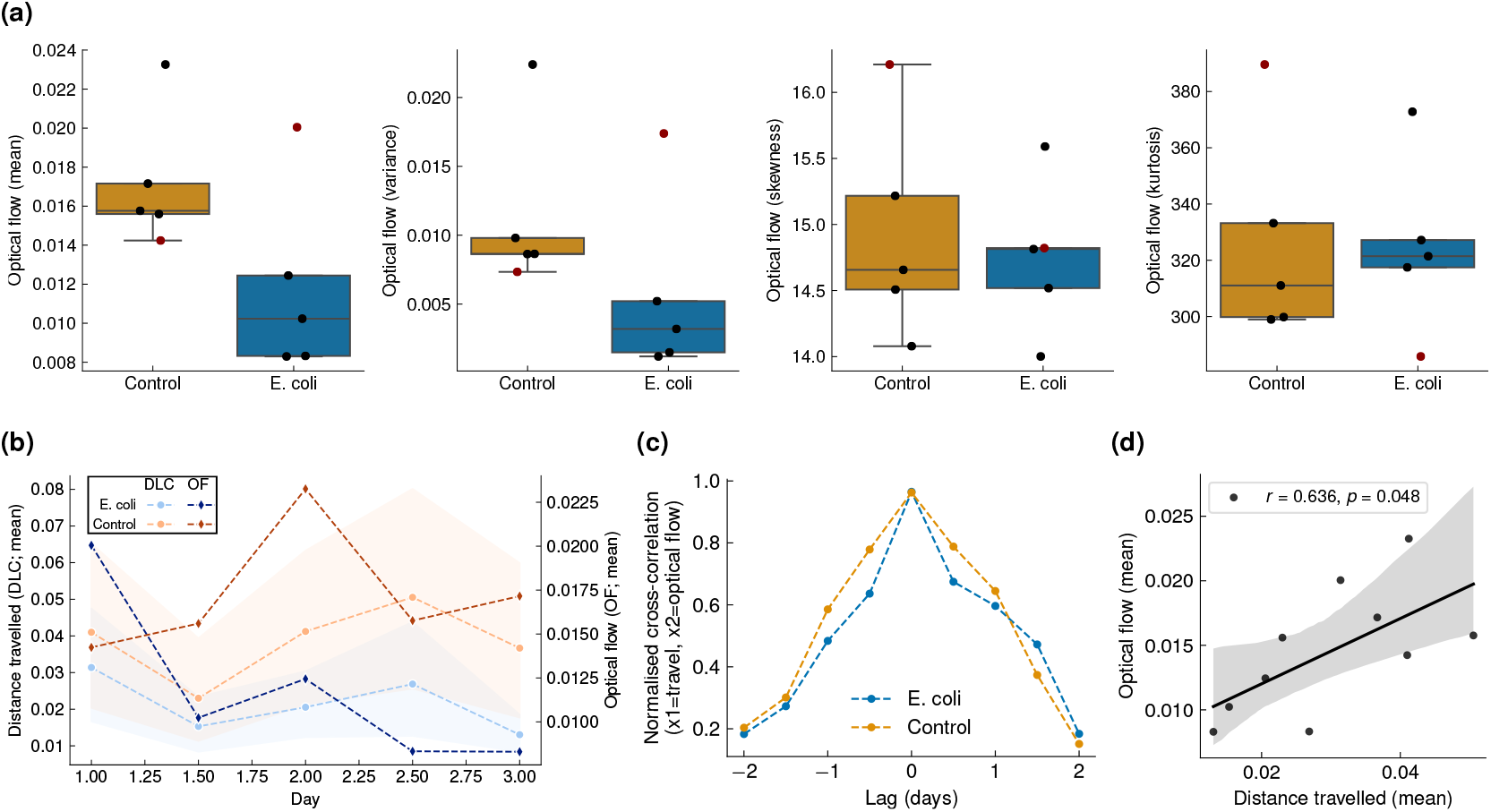
Validation of the travelled distance measured using the multi-animal DeepLabCut (maDLC) [28] against optical flow statistics extracted from a pre-trained SEA-RAFT model [43]. **(a)** Summary statistics (mean, variance, skewness, kurtosis) averaged over each frame and video for each group. Points from the first recording (performed on challenge day) recording are marked in dark red. **(b)** Comparison of the mean optical flow (across frames) against the mean distance travelled (across frames and animals) over time. **(c)** Normalised cross-correlation between the average distance travelled measured using maDLC and the average optical flow. **(d)** Group- and time-independent regression of the mean optical flow (across frames) against the mean distance travelled (across frames and animals). Pearson correlation (*r*): 0.636 (*p* = 0.048). In **(b)** and **(d)**, the shaded areas represent bootstrapped confidence intervals (95%) around the estimates (number of bootstraps = 1,000). In each panel, one point refers to one 20 to 30-min video (see Behavioural data sampling).

## 4. Discussion

In this proof-of-concept study, we demonstrate the capacity of deep learning-based, markerless tracking from video recordings to detect phenotypical differences in broiler chickens following *E. coli* infection. This method holds significant promise for promptly identifying infection or compromised animal well-being at the individual level, without human intervention or intrusive sensors [22] - a capability previously limited to flock-level analyses by conventional techniques [14, 17]. Moreover, the simplicity and objectivity of the statistical analyses derived from video data suggest promising avenues for future research in disease detection and management within the poultry industry. In our analysis, three intuitive features from multi-animal DeepLabCut tracking [28] revealed distinct behavioural patterns between *E. coli* infected and uninfected chickens, corresponding to observed differences in common physiological markers of infection, including serum amyloid A levels, *E. coli* concentrations and organ lesion scores. These features were further validated against optical flow analysis, a traditional computer vision approach that is also promising for measuring animal welfare at industrial scales [12–19].

Both deep learning and optical flow pointed towards greater activity levels in control animals, reflecting a considerable impact of *E. coli* infection on the nervous system. Our data on raised SAA in infected chickens provides evidence that the root cause of behavioural change is likely induced by cytokine-mediated inflammation. Other forms of neurotoxicity that were not explored might also be at play, such as from shiga toxins [52], or gut-brain axis disruption [53]. Previous work has found lower mean optical flow in chickens infected by *Campylobacter* [14], while we observe slight changes in variance, but not in kurtosis after *E. coli* infection. Lower variance in infected individuals is consistent with reduced long-distance movement, while maintained kurtosis indicates stable core short-distance behaviours. Further investigation with larger samples and replicates across different centres will be crucial to confirm the robustness and generalisability of deep learning-based approaches. Additionally, these methods hold significant potential for farms and the poultry industry, particularly for real-time detection of welfare deterioration signs, such as lameness or infections caused by other pathogens.

Identifying infections using videography presents several challenges, some of which are inherent to the livestock industry, where individual animal identification is labour-intensive. Accordingly, we did not preserve bird identity across recordings and therefore did not perform within-subject analyses. Although mixed-effects modelling was used to compare trends between groups, analysing individual differences in behavioural and physiological data would facilitate high-resolution analyses. This warrants future work with more replicates and lifelong video monitoring, or the use of individual features for identification at maximal video resolution. Similarly, we focused on coarse behaviours that might not reflect the behavioural nuances of the immune state of individuals. Future research could investigate a broader spectrum of behavioural responses, e.g., pecking activity, to achieve more precise measurements of feeding behaviour.

Scaling up deep learning-based pose estimation methods to farm-level settings can be achieved via a series of technological and practical solutions. A critical aspect is the ability to perform long-term, continuous monitoring, with the goal of maintaining individual identity over time and potentially detecting early signs of decreased welfare on a per-animal basis. Current tracking of individual body parts becomes challenging when the number of individuals is large (e.g., *>* 50), but solutions are being quickly developed to mitigate the errors arising from crowded environments [54]. Other tracking frameworks, such as TRex [27], allow for tracking up to 256 individuals with reasonable precision, but do not track individual body parts and are still insufficient for poultry farm-level conditions where several thousand animals can coexist. To that end, we posit that a hybrid solution combining a coarse-grained technique, such as optical flow, to detect zones of lower activity with a high-resolution animal tracking framework to identify impaired individuals could constitute a viable solution for curbing outbreaks in farms and thus minimise potential economic losses.

## 5. Conclusion

In this study, we showcase a framework for deep learning-based, markerless tracking of broiler chickens for individual-level detection of behavioural changes associated with *E. coli* infection. The behavioural differences observed in an infected group vs. a control group were validated against physiological markers of infection. More generally, we believe our findings demonstrate the potential of deep learning-based video analysis as an objective tool for outbreak detection in industrial farming, which could be scaled jointly with current flock-level techniques such as optical flow. Livestock settings are complex, and yet advances in data science will continue to disentangle the links between infection, the nervous system, and behaviour for industrial applications. The simplicity and objectivity of the statistical analyses derived from video data, coupled with advancements in deep learning technology, suggest promising avenues for future research in disease outbreak detection and management in livestock.

## 6. Author contributions

N.S., L.L.P., A.M.B., S.B., D.A.D.: study conception and design. L.L.P., N.S.: practical execution. L.L.P., A.M.B.: necropsies and microbiology. N.S.: data analysis. M.P.K., M.I-C., P.R.L.: interpretation of results. L.L.P., A.M.B., D.A.D., S.B.: supervision. N.S., M.P.K., S.B., D.A.D: writing of the original draft. N.S., L.L.P., M.P.K, M.I-C., P.R.L., D.J.L., C.A.D., A.M.B., S.B., D.A.D.: edition of the original draft.

## 9. Acknowledgements

The authors thank Alexandros Katsiferis for valuable insight on analysing the `brms` outputs.

## 10. Data and code availability

All code and data relevant for reproducing the analyses are available at: https://github.com/Neclow/dlc2ecoli. DeepLabCut model weights, training sets, and tracking results are available at: https://doi.org/10.5281/zenodo.15536389.

## A. Appendix

### A.1. Supplementary Figures

**Figure S1.**
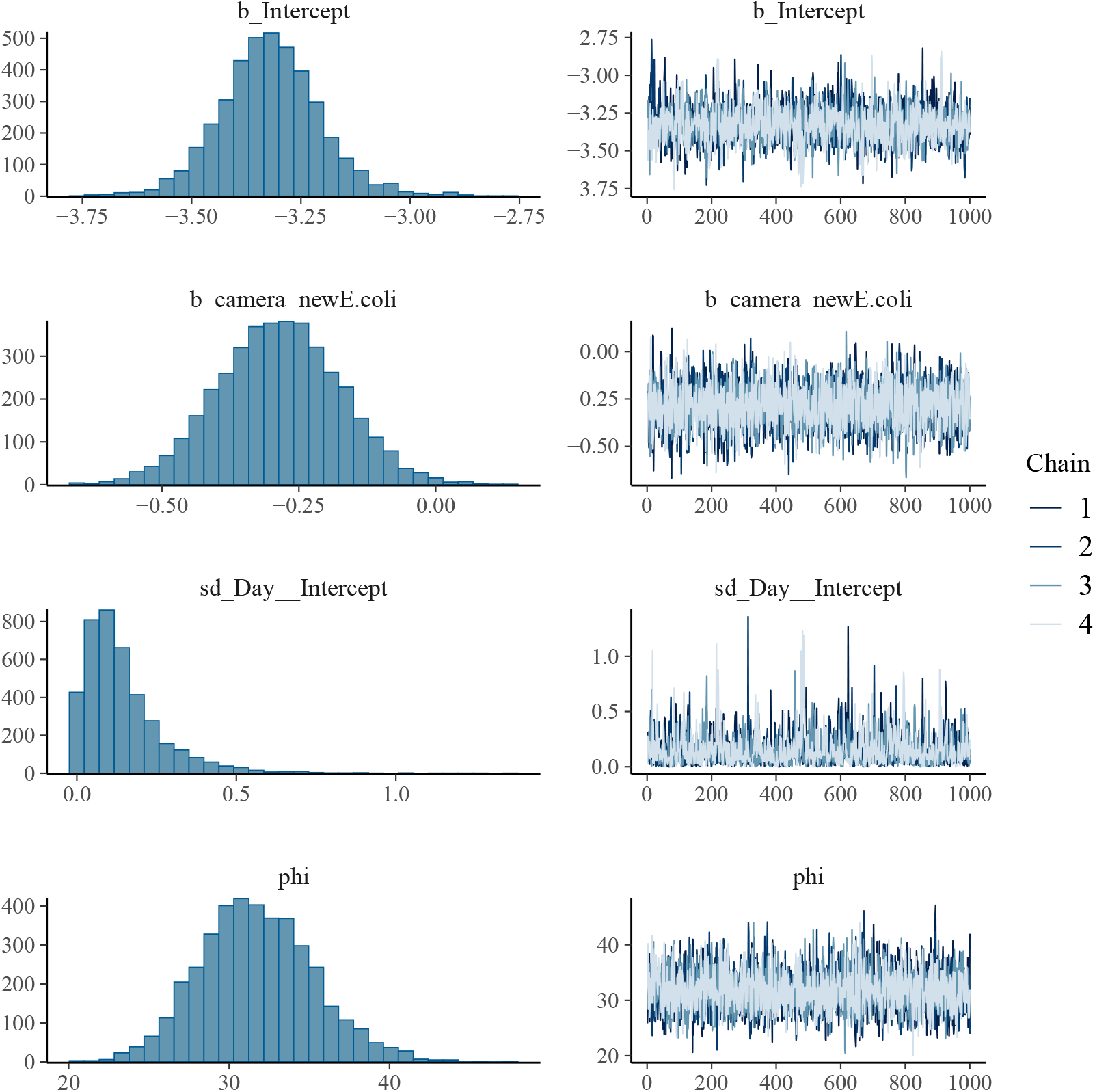
`brms` output for distance travelled, showing density plots for each factor (left) and caterpilar plots (right). Indices: intercept (*b_Intercept*), animal group (between-subject factor; *b_camera_newE*.*coli*), day (within-subject factor; *sd_Day Intercept*), dispersion parameter (*phi*).

**Figure S2.**
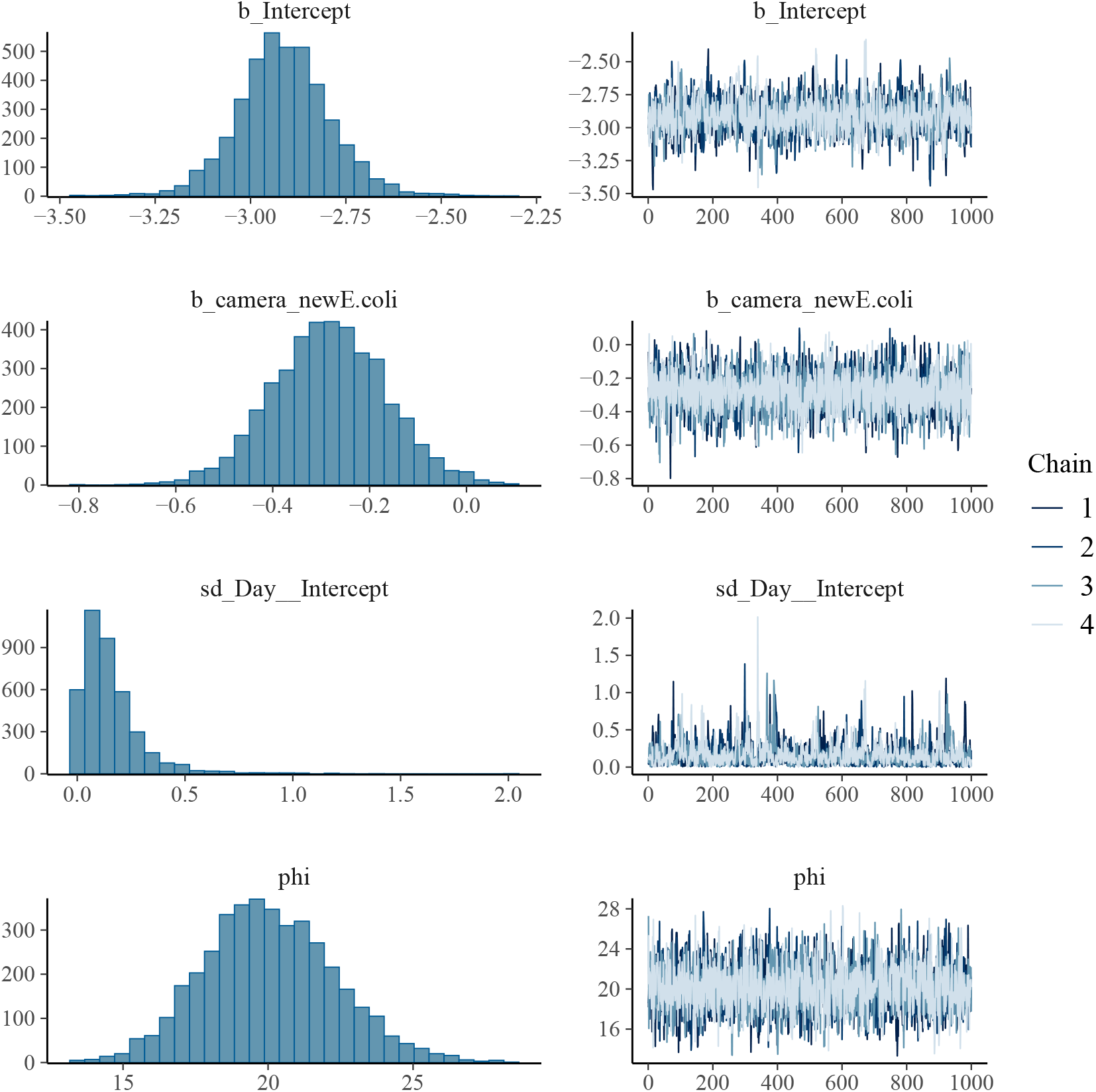
`brms` output for the change in body area, showing density plots for each factor (left) and caterpilar plots (right). Indices: intercept (*b_Intercept*), animal group (between-subject factor; *b_camera_newE*.*coli*), day (within-subject factor; *sd_Day Intercept*), dispersion parameter (*phi*).

**Figure S3.**
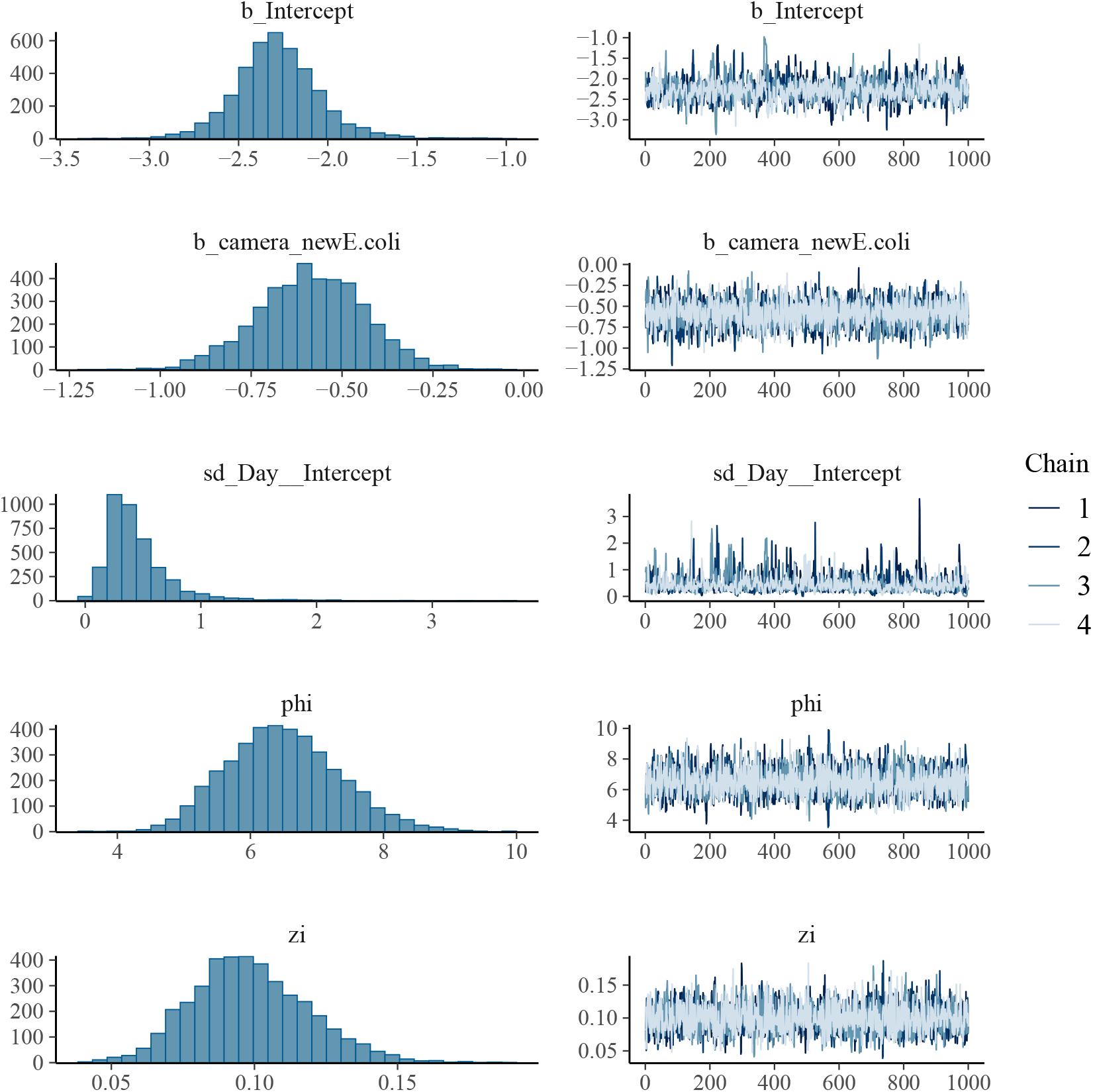
`brms` output for the time spent near the food source, showing density plots for each factor (left) and caterpilar plots (right). Indices: intercept (*b_Intercept*), animal group (between-subject factor; *b_camera_newE*.*coli*), day (within-subject factor; *sd_Day Intercept*), dispersion parameter (*phi*) and zero-inflation parameter (*zi*).

**Figure S4.**
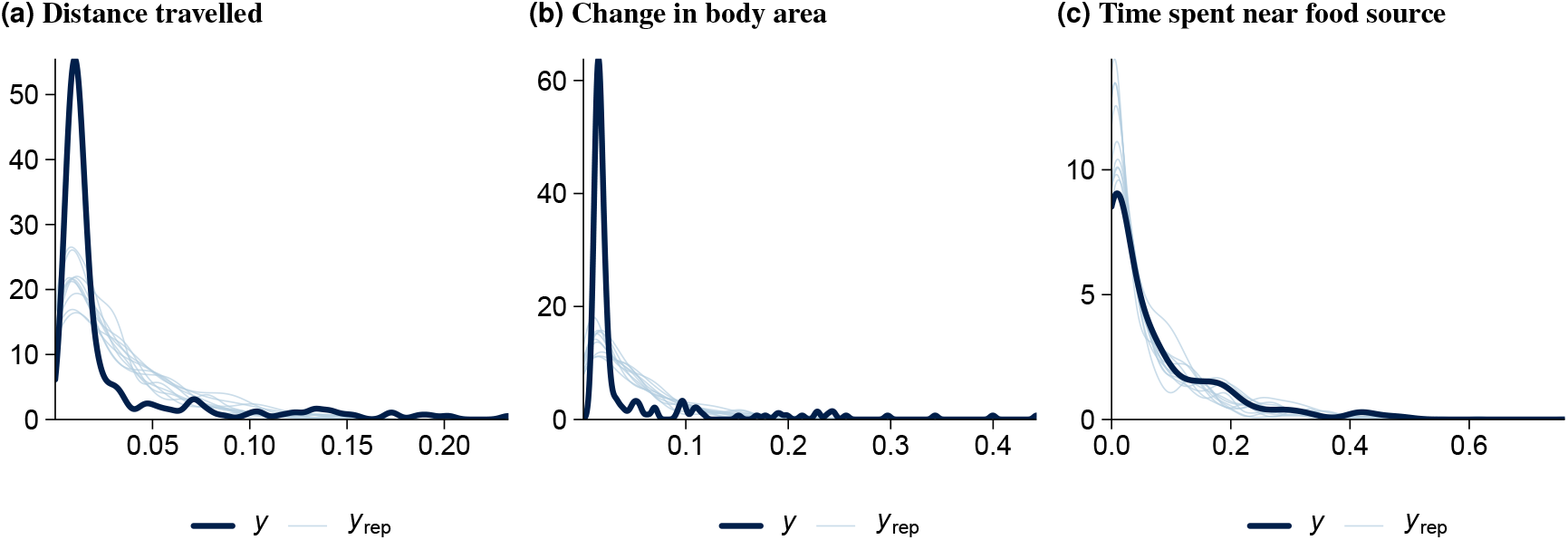
Posterior predictive checks for each model fit. *y*: observed outcome variable, *y*_*rep*_: simulated datasets from the posterior predictive distribution.

### A.2. Supplementary Tables

**Table S1.** Description of lesion score levels.

| Organ | Characteristic | Levels |
| --- | --- | --- |
| **Peritoneum** | Inflammatory reaction | 0: None<br>1: Local at the entrance<br>2: Around the ovary<br>3: In the omentum<br>4: All peritoneum |
|  | Amount of exudate | 0: None<br>1: Sparse<br>2: Some<br>3: Abundant |
|  | Type of exudate | 0: None<br>1: Aqueous<br>2: Lump of pus<br>3: Flakes<br>4: Confluent |
|  | Transparency of peritoneum | 0: Clear<br>1: Nuclear<br>2: Cloudy<br>3: Milky<br>4: Opaque |
| **Liver** | Inflammatory reaction | 0: None<br>1: Fibrin covering < 25%<br>2: Fibrin covering > 25%<br>3: Fibrin covering all the liver |
| **Airsac (left)** | Transparency of airsac | 0: Clear<br>1: Nuclear<br>2: Cloudy<br>3: Milky<br>4: Opaque |
|  | Amount of exudate | 0: None<br>1: Sparse<br>2: Some<br>3: Abundant |
| **Lung (left)** | Amount of exudate | 0: None<br>1: Sparse<br>2: Some<br>3: Abundant |
|  | Congestion | 0: None<br>1: Sparse (< 25% of lung tissue)<br>2: Pronounced (> 25% of lung tissue) |
|  | Lung lesions (oedema) | 0: None<br>1: Slight oedema of the alveolar walls<br>2: Moderate oedematous thickening of alveolar walls with occasional alveoli containing coagulated oedema fluids<br>3: Extensive occurrence of alveolar and interstitial oedema |
|  | Lung lesions (size) | 0: No lesions<br>1: Small lesions between 1 cm<sup>2</sup> and 2 cm<sup>2</sup><br>2: Moderate-size lesions between 3 cm<sup>2</sup> and 4 cm<sup>2</sup><br>3: Extensive lesions with more than 4 cm<sup>2</sup> and covering almost all the area of the lungs |
|  | Granulomas | 0: None<br>1: Few small white caseous/spherical nodules<br>2: Numerous small white caseous/spherical nodules<br>3: Many caseous lesions/nodules and green-gray mold due to sporulation |
| **Spleen** | Proliferation | 0: None<br>1: Little proliferation<br>2: Distinct proliferation |

**Table S2.**
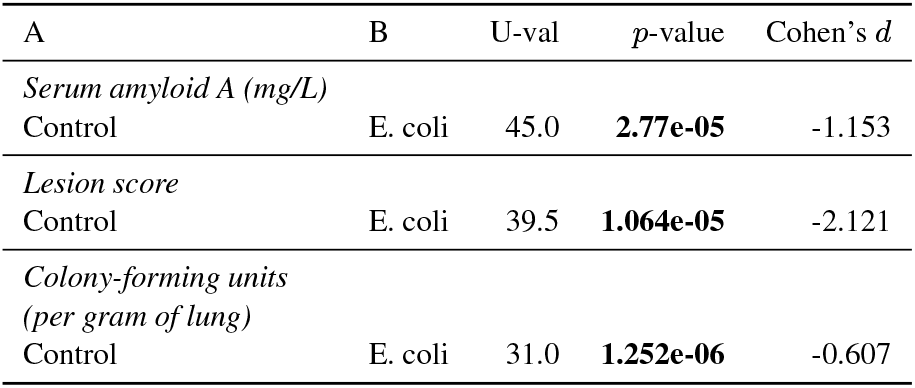
Mann-Whitney U tests to assess between-group differences in serum amyloid A levels, lesions scores, and number of colony-forming units.

**Table S3.** `brms` summary tables for the behavioural features. Reference group: Control (uninfected). Significant differences (determined if the confidence intervals exclude zero) with respect to the negative control are shown in bold. CI: confidence interval (l: lower, u: upper). 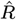: Gelman-Rubin statistic for Markov chain Monte Carlo convergence; convergence is deemed suitable when 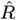 is close to 1. ESS: effective sample size.

| | Estimate | Est.Error | l-95% CI | u-95% CI | $\hat{R}$ | Bulk ESS | Tail ESS |
| --- | --- | --- | --- | --- | --- | --- | --- |
| *Distance travelled* |  |  |  |  |  |  |  |
| Intercept | -3.32 | 0.116 | -3.543 | -3.079 | 1.003 | 2353.594 | 1771.796 |
| E. coli | -0.286 | 0.112 | **-0.506** | **-0.069** | 1.002 | 3717.03 | 2639.991 |
| *Rate of change in body area* |  |  |  |  |  |  |  |
| Intercept | -2.912 | 0.125 | -3.148 | -2.658 | 1.001 | 1750.502 | 1543.872 |
| E. coli | -0.283 | 0.117 | **-0.518** | **-0.049** | 1.001 | 3034.374 | 2758.279 |
| *% Time near food source* |  |  |  |  |  |  |  |
| Intercept | -2.281 | 0.245 | -2.74 | -1.761 | 1.002 | 1333.479 | 1306.324 |
| E. coli | -0.585 | 0.148 | **-0.883** | **-0.302** | 1.001 | 3112.788 | 2167.955 |

**Table S4.**
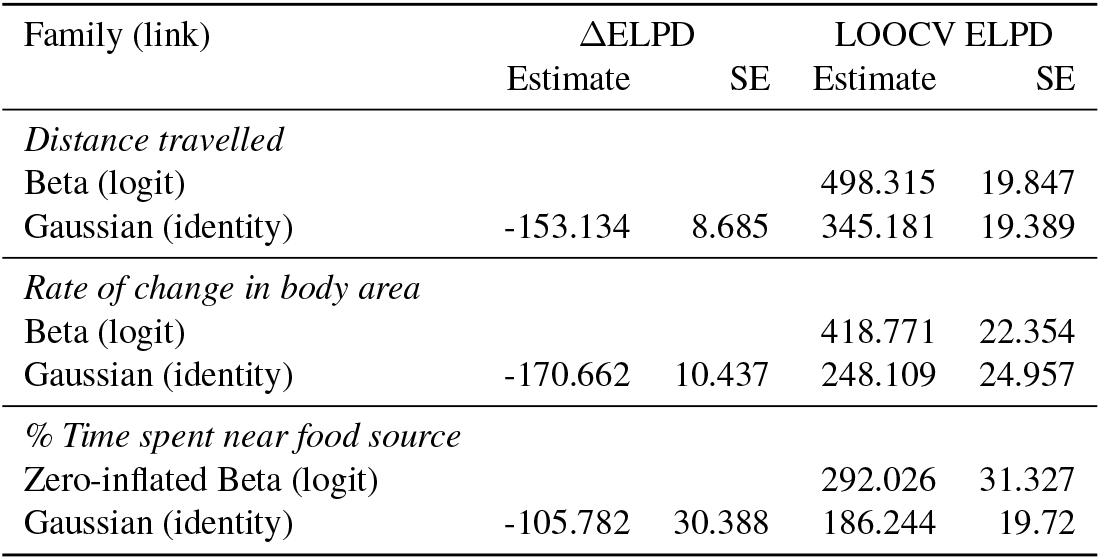
Comparison of different families for mixed-effects modelling using brms. ELPD: expected log pointwise predictive density. LOOCV: leave-one-out cross-validation. SE: standard error.

| Family (link) | ΔELPD | | LOOCV ELPD | |
| --- | --- | --- | --- | --- |
|  | Estimate | SE | Estimate | SE |
| *Distance travelled* |  |  |  |  |
| Beta (logit) |  |  | 498.315 | 19.847 |
| Gaussian (identity) | -153.134 | 8.685 | 345.181 | 19.389 |
| *Rate of change in body area* |  |  |  |  |
| Beta (logit) |  |  | 418.771 | 22.354 |
| Gaussian (identity) | -170.662 | 10.437 | 248.109 | 24.957 |
| *% Time spent near food source* |  |  |  |  |
| Zero-inflated Beta (logit) |  |  | 292.026 | 31.327 |
| Gaussian (identity) | -105.782 | 30.388 | 186.244 | 19.72 |

